# Microevolutionary system identification and climate response predictions

**DOI:** 10.1101/2022.08.15.503935

**Authors:** Rolf Ergon

**Affiliations:** University of South-Eastern Norway

**Keywords:** Climate response predictions, Microevolutionary system identification, MIMO system, Prediction error method, Reaction norm model, Reference environment

## Abstract

Microevolutionary system identification was introduced in Ergon (2022), with the specific purpose to show that predictions of genetic adaptations to climate change require that environmental reference values are properly defined. The theoretical development was then limited to single-input single-output (SISO) systems, and the simulations used a toy example with spring temperature as input and mean breeding date as output. Generations were assumed to be non-overlapping. Here, the theory is extended to cover multiple-input multiple-output (MIMO) systems, while the simulation example uses two environmental inputs (spring temperature and rainfall) and two adaptive phenotypic outputs (breeding date and breeding habitat). These extended simulations reveal difficulties involved in predictions of genetic adaptations for complex systems based on short data, where the reference environment values are not included.

## 1 Introduction

Wild populations respond to changing environments by means of phenotypic plasticity and microevolution (Ergon, 2019), and especially climate change responses have been extensively studied. The aim is then to disentangle phenotypic changes owing to genetically based microevolution, caused by natural selection, and changes due to individual phenotypic plasticity. Microevolution here refers to genetical changes that occur over time in a specific population under influence of the environment. Such changes may be relatively fast (in evolutionary terms), compared to the changes caused by macroevolution, which also involves interaction between several and often very different populations, possibly also the formation of new and distinct species.

Relying on 11 review articles, including reviews of altogether 66 field studies, Merilä and Hendry (2014) arrived at the conclusion that evidence for genetic adaptation to climate change has been found in some systems, but that such evidence is relatively scarce. They also concluded that more studies were needed, and that these must employ better inferential methods.

Ergon (2022) focused on a problem that appears to be overlooked in the field studies reviewed in Merilä and Hendry (2014), and in the experimental quantitative genetics community in general (Shaw and Etterson, 2012). It is obvious that for all evolutionary systems with interval-scaled environmental variables *u_t_*, as for example temperature in °C, a suitable zero-point (reference environment) *u_ref_* must be chosen, and this should not be done arbitrarily. As discussed in Ergon (2022), the proper zero-point is the environment where the expected geometric mean fitness has a global maximum, and thus the environment the population is fully adapted to. Fitness is here a measure of reproductive success, for example the number of offspring.

Climate response studies are based on input-output data that primarily are collected in field studies of wild populations of animals, plants and other organisms, for example mating date as function of spring temperature. From an engineering control point of view, such data quite naturally point towards use of system identification methods, and Ergon (2022) therefore introduced microevolutionary system identification as a tool in the field of experimental quantitative genetics.

As a theoretical study system, Ergon (2022) used the intercept-slope individual reaction norm model (Fig.2, Ergon, 2019; Fig.1, Ergon, 2022)

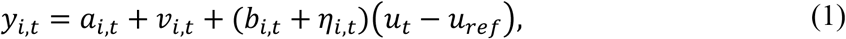

where *u_t_* – *u_ref_* and *y_i,t_* are the environmental cue and the individual phenotypic value, respectively, as functions of time *t* measured in generations. Here, *a_i,t_* and *b_i,t_* are the additive genetic components of the intercept and plasticity slope, respectively, while *v_i,t_* and *η_i,t_* are independent and identically distributed (iid) zero mean normal non-additive effects. As done in Lande (2009) and Ergon and Ergon (2017), we may consider the individual reaction norm intercept *a_i,t_* + *v_i,t_*, and the individual plasticity slope *b_i,t_* + *η_i,t_*, as two quantitative traits in their own rights. Microevolution thus involves changes in the population mean trait values 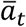 and 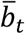 from generation to generation. The generations are here assumed to be non-overlapping.

**Figure 1.**
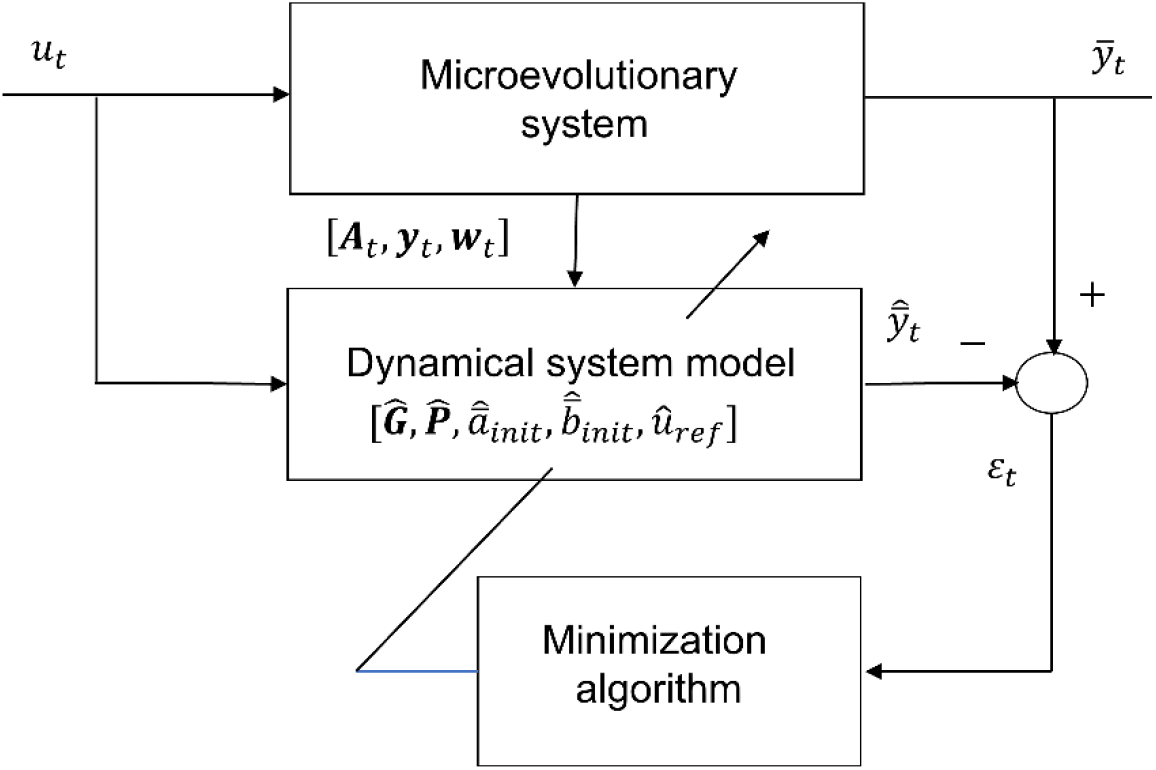
Block diagram of microevolutionary PEM for SISO system, with dynamical tuning model based on an intercept-slope reaction norm model with mean traits 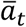 and 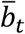. Here, *u_t_* and 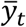 are the known environmental input and the known mean phenotypic value at time *t*, respectively. ***A_t_*** is the additive genetic relationship matrix, which here is assumed to be ***A_t_*** = ***I***, while ***y_t_*** and ***w_t_*** are vectors of individual phenotypic and relative fitness values, respectively. The 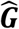 and 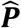 matrices include the system parameters, while 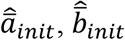 and 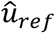 are the initial mean trait values and the reference environment, respectively. Assuming data over *T* generations, all these model parameters are tuned until 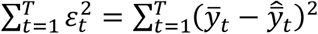 is minimized, with 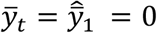 and 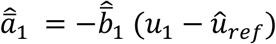.

**Figure 2.**
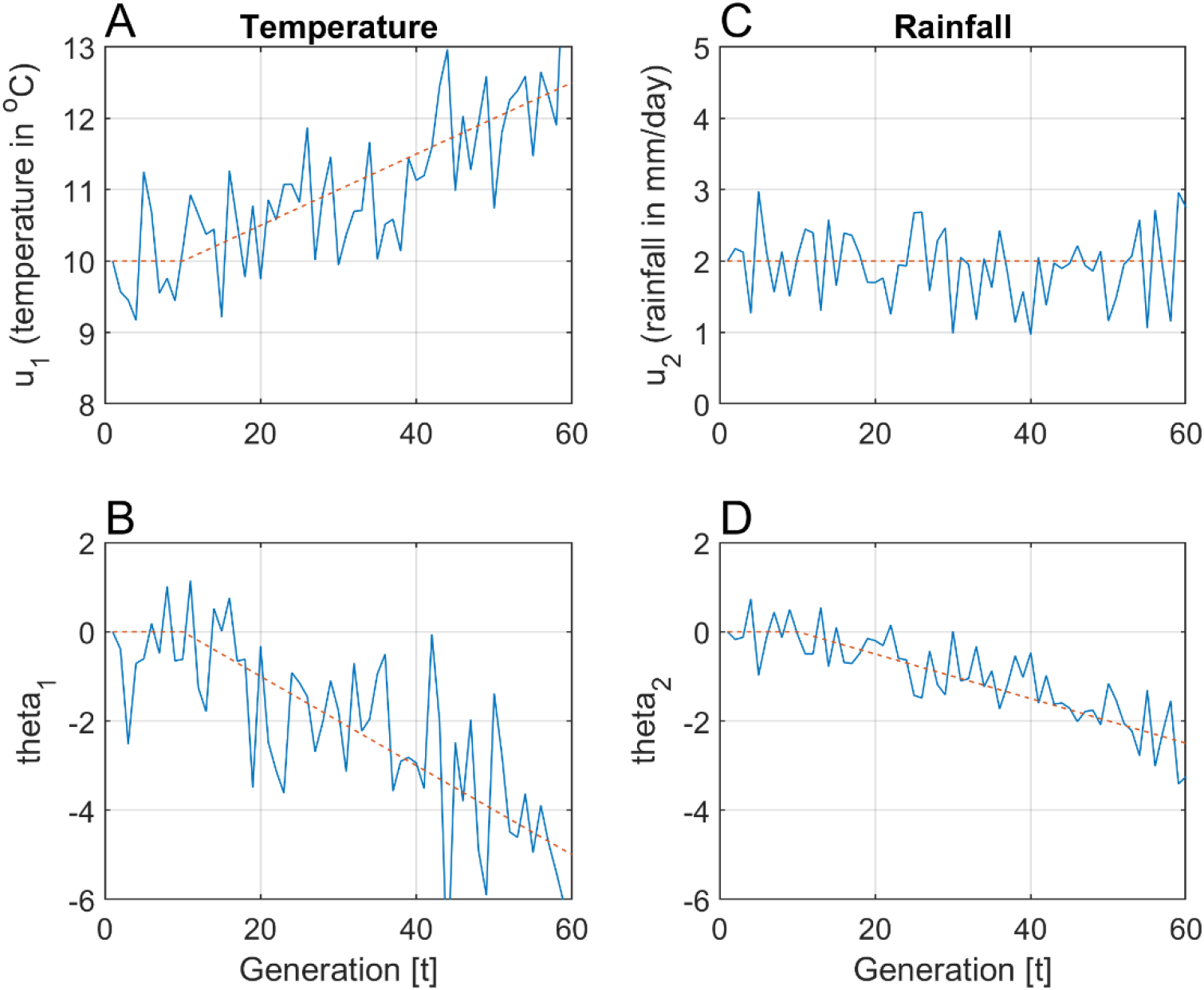
Typical input signals for simulation example, with three noisy ramp functions starting at generation *t* = 10 (1970). Here, *μ*_*U*_1_,*t*_ is a ramp function and *μ*_*U*_2_,*t*_ = 2 (dashed magenta lines), while *μ*_Θ_1_,*t*_ = −2(*μ*_*U*_1_,*t*_ – 10) and *μ*_Θ_2_,*t*_ = −(*μ*_*U*_1_,*t*_ – 10) (dashed magenta lines). Other numerical values are 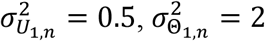 and 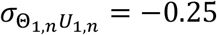 (as in Ergon, 2022), and 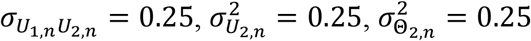 and 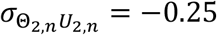.

From Equation (1) follows the mean trait reaction norm model 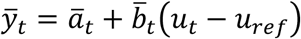, and from this simple equation follows the basic questions discussed in Ergon (2022). How can *u_ref_* be estimated, and how can the evolution of 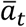 and 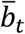 be predicted, provided that *u_t_* and *y_i,t_* are known? And how will the predictions be affected by errors in the estimated or assumed value 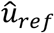? It turns out that in order to answer these questions we also need information on individual fitness values *W_i,t_*. The discussion in Ergon (2022) was limited to single-input single-output (SISO) systems, where parameter estimates, and mean trait predictions were found by use of a prediction error method (PEM) (Ljung, 2002). Here, the discussion is extended to multiple-input multiple-output (MIMO) systems, which further reveals the difficulties involved in the disentanglement of plasticity and microevolutionary effects on phenotypic change.

The intercept and plasticity slope traits in Equation (1) are characterized by the additive genetic covariance matrix

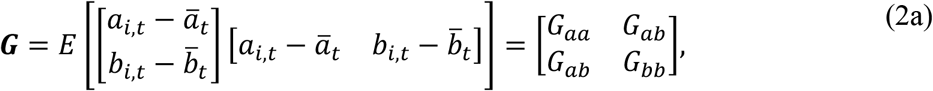

and the phenotypic covariance matrix

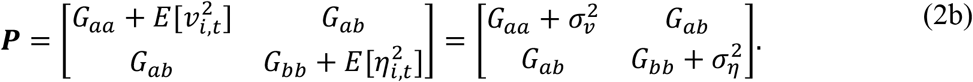

These matrices will in general be functions of time, but for simplicity they are here assumed to be constant, as is often done in theoretical work (e.g., Lande, 2009).

In order to predict the evolution of 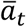 and 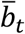 over one or several generations, we must estimate the parameters in ***G*** and ***P***. We must also find initial values 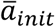 and 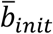, and the environmental reference value *u_ref_* must be either estimated or assumed known. As proposed in Ergon (2022), all of this can be achieved by use of a prediction error method (PEM) as shown in Fig 1. This is an output error model (Ljung, 2002), where the output 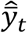 from a tuning model is directly compared with the true output 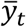. Fig. 1 is general in the sense that there may exist a relationship matrix ***A**_t_* that is not a unity matrix, i.e., that there may be inbreeding in the population. However, Ergon (2022) assumed the special case with ***A_t_* = *I***, which means that breeding is based on random mating in a large population. This is assumed also here.

A main problem in the PEM in Fig. 1 is to find how the output 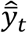 from the tuning model is determined by the predicted mean traits 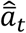 and 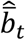. In Ergon (2022) this theoretical problem was solved by use of a linear transformation of the vector 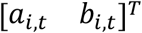 onto the vector 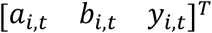. Here, this solution is extended from the SISO case to cover also MIMO systems, with obvious changes in the notation in Fig. 1.

The main practical contribution in Ergon (2022) was to point out the necessity of a properly chosen environmental reference value, and to show that errors in 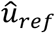 lead to errors in predicted changes in 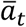 and 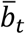 over time. The background for the simulations was the fact that the global mean temperature since 1970 has increased as a noisy ramp function with around 0.017 °C/*year*, and that it up to 1970 was fairly constant (NASA, 2019). The simulations therefore assumed a population that was fully adapted to the temperature before 1970, which was thus used as the temperature reference, but they also assumed that input-output data was available only for the last 30 years.

In Ergon (2022), the use of a reference environment that was not within the range of input data used in the SISO system identification algorithm did not create a problem. However, the MIMO simulations in the present article show that correct reference values 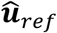 outside the range of input data, may lead to severe convergence problems in the PEM method according to Fig. 1. The only practical solution may thus be to use the first values ***u***_1_ in the multivariate input data series (or values close to that) as reference values, and thus accept that this leads to prediction errors. There is however a certain possibility for correction of these errors, provided that the reference environment is approximately known.

## 2 Theory

### 2.1 Prediction equations

As shown in Ergon (2022), mean trait predictions based on the reaction norm model (1) are found from

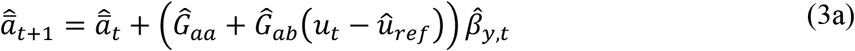

and

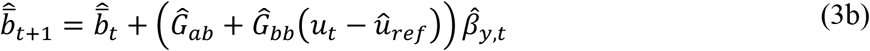

where 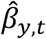 is the estimated selection gradient (Lande, 2009),

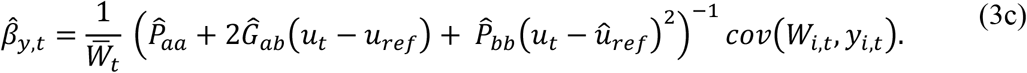

Note that these equations are valid only when the genetic relationship matrix is a unity matrix (Ch. 26, Lynch & Walsh, 1998).

For a MIMO system with two input signals and two output signals, the individual reaction norm model (1) is replaced by

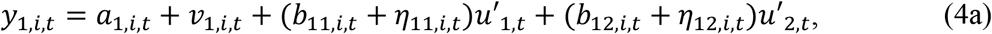

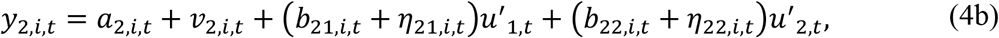

where 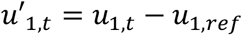 and 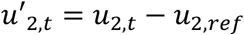. Here, all traits may be correlated, such that for example 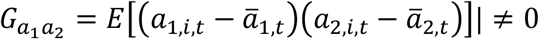.

More compact and general for *r* input signals and *m* output signals we get

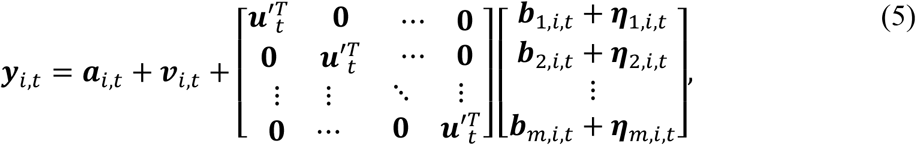

where ***y**_i,t_*, ***a**_i,t_* and ***v**_i,t_* are *m* × 1 vectors, while 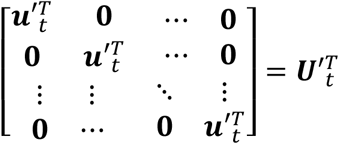 is an *m* × *rm* input signal matrix, and 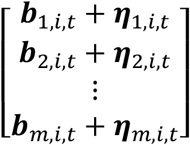 an *rm* × 1 vector. Here 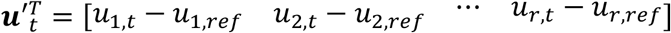 and 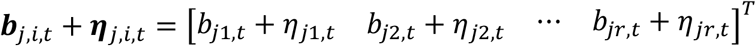.

The total covariance matrices for the system (5) are 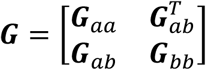, and 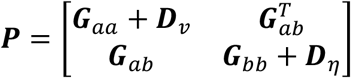, where ***G**_aa_*, ***G**_ab_* and ***G**_bb_* are *m* × *m*, *rm* × *m* and *rm* × *rm* matrices, respectively, while (in MATLAB notation)

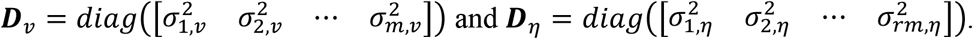

A comparison with Equations (3a,b) now leads to the generalized prediction equations

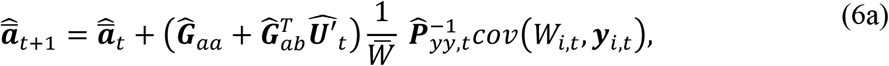

and

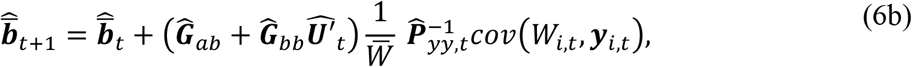

where 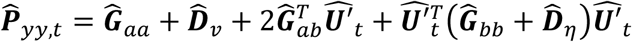. Note that we here may set for example 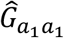 to any value, and estimate the rest of the parameters in relation to this value (because 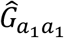 appears in 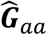 and 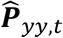).

With given initial values, Equations (6a,b) give 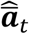 and 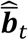, and thus

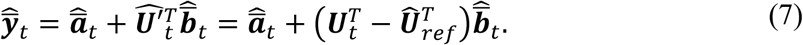

From the tuning model in Fig. 1 we thus find predictions 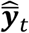 based on 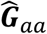 etc., and what remains is to minimize a criterion function

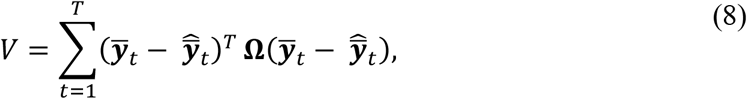

where **Ω** is a diagonal weighting function, possibly **Ω** = ***I**_m_*.

### 2.2 Prediction errors caused by errors in *u_ref_*

Prediction errors caused by errors in the assumed reference environment 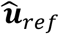 can be found as for the SISO case in Ergon (2022), only somewhat more complicated. With a reference environment 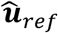 instead of ***u**_ref_*, and thus an input signal matrix 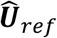 instead of ***U**_ref_*, predictions based on Equation (7) can be written as

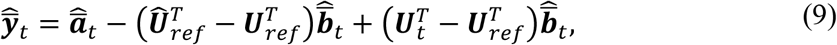

where 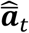 and 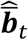 are found from Equations (6a,b), assuming initial values known.

For small parameter values in ***P**_bb_*, i.e., when ***P**_bb_ →* 0 and ***G**_ab_* → 0, it follows from Equation (6a) that 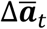 is independent of ***u**_ref_*, and that 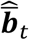 is constant. This results in 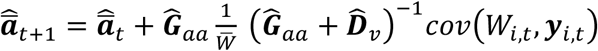, such that only 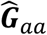 (except for 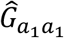) and 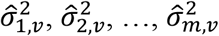 must be tuned in order to minimize 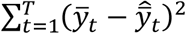. In this case an error in 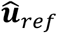 has very little effect on the change in 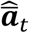 per generation.

For larger values of ***P**_bb_*, the predicted changes per generation, 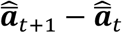, will be affected by an error in 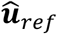, and with ***G**_ab_ =* 0 good predictions 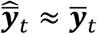 for *t* = 1 to *T* can then only be obtained by parameter tuning such that 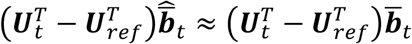 over all generations. That is possible because ***u**_ref_* in Equation (6b) appears in both 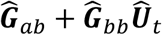 and 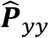. According to Equation (9) we then find 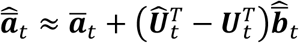, which as shown in Section 3 may result in large errors in predicted changes of 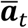 over time. This corresponds to the result in the SISO case discussed in Ergon (2022), except that we in the MIMO case cannot expect good predictions of the elements in 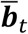, but only of the product 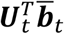. Note that if ***u**_ref_* is (approximately) known, it is possible to find a corrected estimate 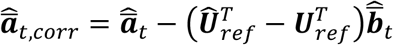.

## 3 Simulation results

### 3.1 Description of toy example

In the toy example in Ergon (2022), the environmental input was a noisy positive trend in spring temperature, resulting in a noisy negative trend in mean breeding (clutch-initiation) date for a certain bird species, approximately as in Fig. 2 in Bowers et al. (2016). The individual phenotypic values were discrete, with days as unit. The individual (mid-parent) fitness values were integers from 0 to 10, with number of offspring as unit. Generations were assumed to be non-overlapping, and the population size was assumed to be constant.

Here, the example is extended to include a second input variable with a constant mean value, and with variations from year to year that are somewhat correlated with the variations in spring temperature. This input may for example be a measure of rainfall. The example also includes a second adaptive phenotype, that might be the breeding habitat, as discussed in Chalfoun and Schmidt (2012). We thus have a MIMO system, with two input signals and two output signals, as in Equations (4a,b).

In the new simulations the two environmental reference values are assumed to be known from historical data, i.e., it is assumed that the population was fully adapted to the stationary stochastic environment before the onset of anthropogenic and global climate change around 1970. The essential question is then how well microevolutionary changes in mean intercepts and plasticity slopes can be predicted by means of the PEM method in Fig. 1. Two cases will be tested, one with long input-output data that include the reference environments, and one with short data that do not include the reference environments.

### 3.2 True model, fitness function, and environmental input signals

Assume a true system with individual reaction norm model according to Equations (4a,b), which by use of the multivariate breeder’s equation (Lande, 1979) gives the state-space model

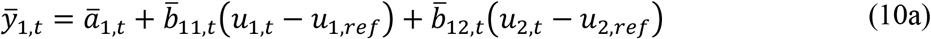

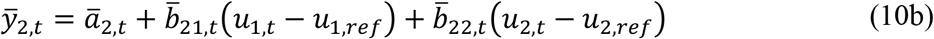

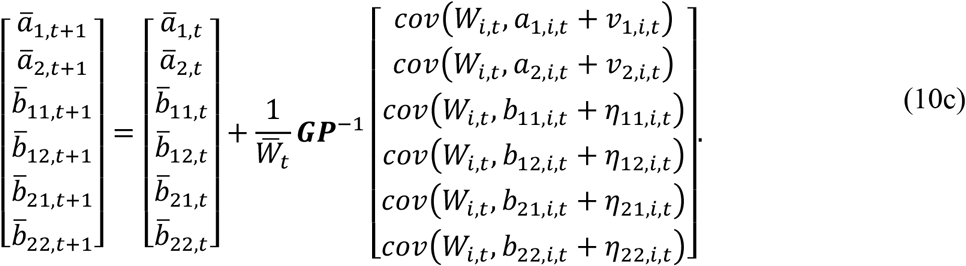

Also assume the additive genetic and phenotypic covariance matrices

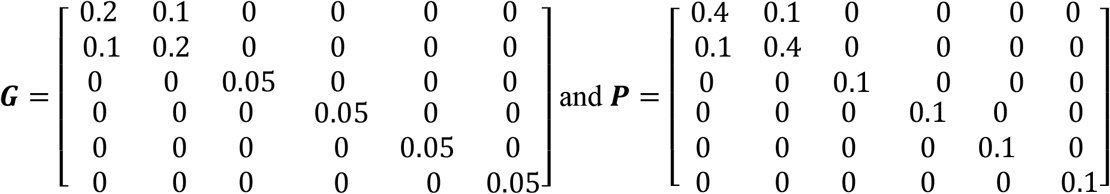

The individual fitness function is assumed to be rounded values of

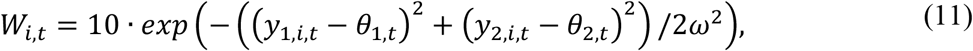

where *θ*_1,*t*_ and *θ*_2,*t*_ are the phenotypic values that maximize fitness, while *ω*^2^ = 10. The discrete values of *W_i,t_* (number of offspring) are thus integers from 0 to 10.

As in Ergon (2022) we assume a stationary or slowly varying mean value *μ*_*U*_1_,*t*_ of a stochastic environment (spring temperature), with added iid zero mean normal random variations 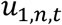 with variance 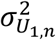, i.e., *u*_1,t_ = *μ*_*U*_1_,*t*_ + *u*_1,*n*,*t*_. Also assume a constant mean value *μ*_*U*_2_,*t*_ of a second stochastic environment (rainfall), with added iid zero mean normal random variations *u*_2,*n,t*_ with variance 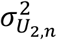, i.e., *u*_2,t_ = *μ*_*U*_2_,*t*_ + *u*_2,*n*,*t*_, and that *u*_1,*n,t*_ and *u*_2,*n,t*_ are correlated with covariance *σ*_*U*_1,*n*_ *U*_2,*n*__. In a corresponding way assume that 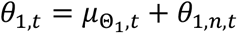, where *θ*_1*n,t*_ is iid zero mean normal with variance 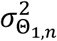, and where *u*_1,*n,t*_ and *θ*_1,*n,t*_ are correlated with covariance 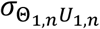, as described in more detail in Ergon (2022). Further assume that *θ*_2,*t*_ = *μ*_Θ_2_,*t*_ + *θ*_2,*n,t*_, where *θ*_2*n,t*_ is iid zero mean normal with variance 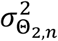 and where *u*_2,*n,t*_ and *θ*_2,*n,t*_ are correlated with covariance *σ*_Θ_2,*n*_*U*_2,*n*__. Finally assume that the population is fully adapted to a stationary stochastic environment with *μ*_*U*_1__ = *u*_1,*ref*_ = 10 °C (as in Fig. 1 in Ergon, 2022)), *μ*_*U*_2__ = 2 (the zero-point for rainfall), and *μ*_Θ_1__ = *μ*_Θ_2__ = 0. Data were generated for 60 generations, with typical input data as shown in Fig. 2 (as mean values in breeding season).

### 3.3 Model choice

The choice of tuning model in Fig. 1 can be guided by how well the model output signals 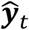 follow the true outputs ***y**_t_*. For the true system in Equations (10a,b,c) and (11), this is illustrated in Fig. 3.

**Figure 3.**
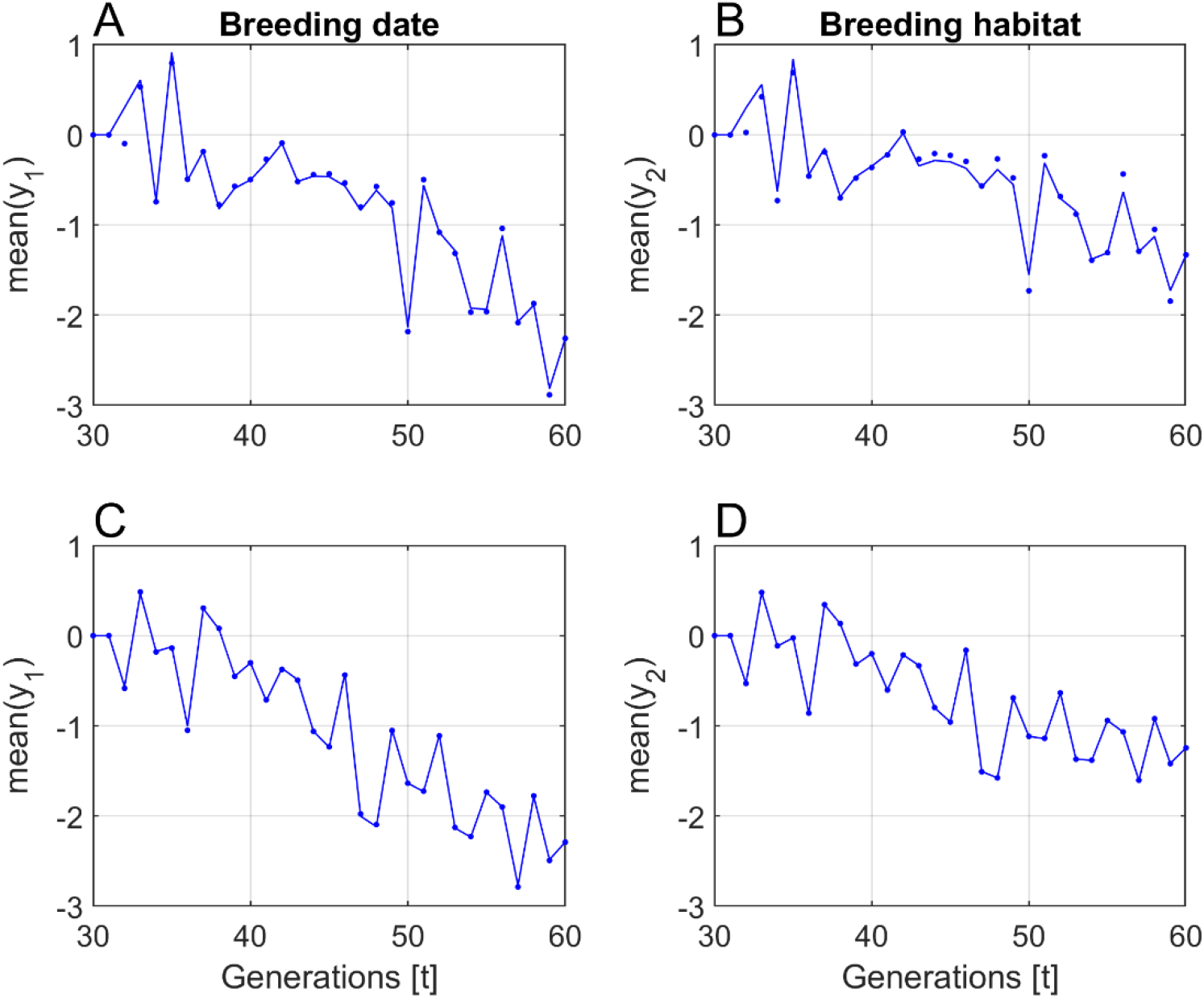
True outputs (solid lines) and outputs from tuning model (dots), for two choices of tuning model, and with use of input-output data from the last 30 generations in Fig. 2. The responses in panels A and B are obtained with the true reference environments *u*_1,*ref*_ = *u*_1,1_ = 10 and *u*_2,*ref*_ = *u*_2,1_ = 2, which leads to convergence problems. The responses in panels C and D are obtained with assumed reference environments *u*_1,*ref*_ = *u*_1,31_ and *u*_2,*ref*_ = *u*_2,31_, which in most realizations gives good convergence, but prediction errors as discussed in Subsection 2.2. These errors can to some degree be corrected if the true values of *u*_1,*ref*_ and *u*_2,*ref*_ are known.

### 3.4 Parameter estimation and mean trait prediction results

Parameter estimates were found by use of the MATLAB function *fmincon* in the PEM method in Fig. 1. The criterion function (8) was 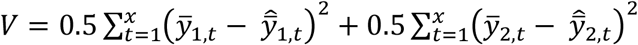, where *x* was either 60 (Case 1) or 30 (Case 2).

Given the model in Equations (10a,b.c) and (11), there are in all 16 parameter values to be estimated (while 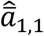 and 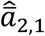 follow from Equations (10a,b) with 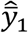 and 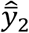 set to zero). In the optimizations, the initial values of 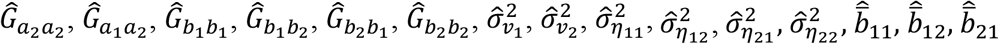 and 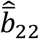 were set to zero. The true value 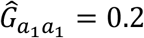 was used, such that estimates of other ***G*** and ***P*** parameters are found relative to that value.

Results with use of input-output data from *t* = 1 to 60 (*x* = 60) with population size *N* = 400 are given in Table 1 (Case 1), but the table also includes results when only data from *t* = 31 to 60 (*x* = 30) were used (Case 2). Note that Case 2 does not make use of data from generations before the start of the ramp functions, where the population is assumed to be fully adapted to the environment. This is the most realistic case, because field data rarely go as far back as before 1970. In both cases, the first available input values were used as environmental reference values, i.e., either 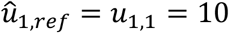 and 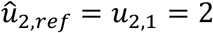 (Case 1), or 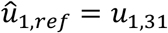 and 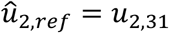 (Case 2). Results are presented as mean values and standard errors, *Mean* ± *SE*, based on 100 repeated simulations with different realizations of random inputs. Note that a modified Case 2 with use of the true reference environments 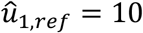 and 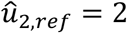 for almost every realization result in convergence problems (Fig. 3, panels A and B).

**Table 1.** Parameter estimation results with true system responses generated by means of Equations (10a,b,c) and (11). Results are for cases with population size *N* = 400 and perfect observations of *y*_1,*i,t*_, *y*_2,*i,t*_, and *W_i,t_*, and they are based on 100 simulations with different realizations of all random input variables. Case 1: *x* = 60, 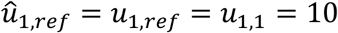 and 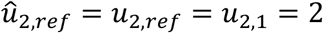. Case 2: 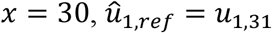 and 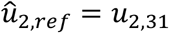. Here, 14% of the simulations were discarded because the final value of the criterion function was *V >* 0.025, which approximately is the limit where it is possible to detect poor convergence by inspection of plots as in Fig. 3.

| Parameter<br>etc. | True<br>value | Results<br>Case 1 | Results<br>Case 2 |
| --- | --- | --- | --- |
| $\hat{G}_{a_2 a_2}$ | 0.2 | $0.2024 \pm 0.0297$ | $0.1954 \pm 0.1328$ |
| $\hat{G}_{a_1 a_2}$ | 0,1 | $0.0997 \pm 0.0034$ | $0.0758 \pm 0.0219$ |
| $\hat{G}_{b_1 b_1}$ | 0.05 | $0.0500 \pm 0.0023$ | $0.0779 \pm 0.0464$ |
| $\hat{G}_{b_1 b_2}$ | 0.05 | $0.0494 \pm 0.0108$ | $0.0158 \pm 0.0317$ |
| $\hat{G}_{b_2 b_1}$ | 0.05 | $0.0502 \pm 0.0123$ | $0.0624 \pm 0.0595$ |
| $\hat{G}_{b_2 b_2}$ | 0.05 | $0.0496 \pm 0.0170$ | $0.0413 \pm 0.0527$ |
| $\hat{\sigma}_{v_1}^2$ | 0,2 | $0.2008 \pm 0.0081$ | $0.1752 \pm 0.0549$ |
| $\hat{\sigma}_{v_2}^2$ | 0,2 | $0.2045 \pm 0.0509$ | $0.1654 \pm 0.1754$ |
| $\hat{\sigma}_{\eta_{11}}^2$ | 0.05 | $0.0500 \pm 0.0036$ | $0.0819 \pm 0.0536$ |
| $\hat{\sigma}_{\eta_{12}}^2$ | 0.05 | $0.0504 \pm 0.0261$ | $0.0459 \pm 0.0596$ |
| $\hat{\sigma}_{\eta_{21}}^2$ | 0.05 | $0.0492 \pm 0.0169$ | $0.0689 \pm 0.0629$ |
| $\hat{\sigma}_{\eta_{22}}^2$ | 0.05 | $0.0646 \pm 0.0527$ | $0.0536 \pm 0.0671$ |
| $\hat{b}_{11}$ | — | $-0.5004 \pm 0.0034$ | $-0.5897 \pm 0.0589$ |
| $\hat{b}_{12}$ | — | $0.0004 \pm 0.0037$ | $0.0035 \pm 0.0342$ |
| $\hat{b}_{21}$ | — | $-0.4996 \pm 0.0029$ | $-0.5247 \pm 0.0154$ |
| $\hat{b}_{22}$ | — | $-0.0003 \pm 0.0031$ | $-0.0210 \pm 0.0147$ |

Table 2 shows the relative errors in total change of mean trait predictions over *x* generations, computed as 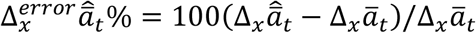 etc., where 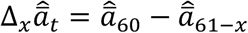 and 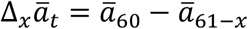. Here, *x* is either 60 (Case 1) or 30 (Case 2). As discussed in Subsection 2.2, Table 2 also includes results for corrected estimates 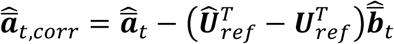 under the assumption that *u*_1,*ref*_ = 10 and *u*_2,*ref*_ = 2 are known. Also here, the results are presented as mean values and standard errors, *Mean* ± *SE*, based on the same repeated simulations as used for Table 1.

**Table 2.**
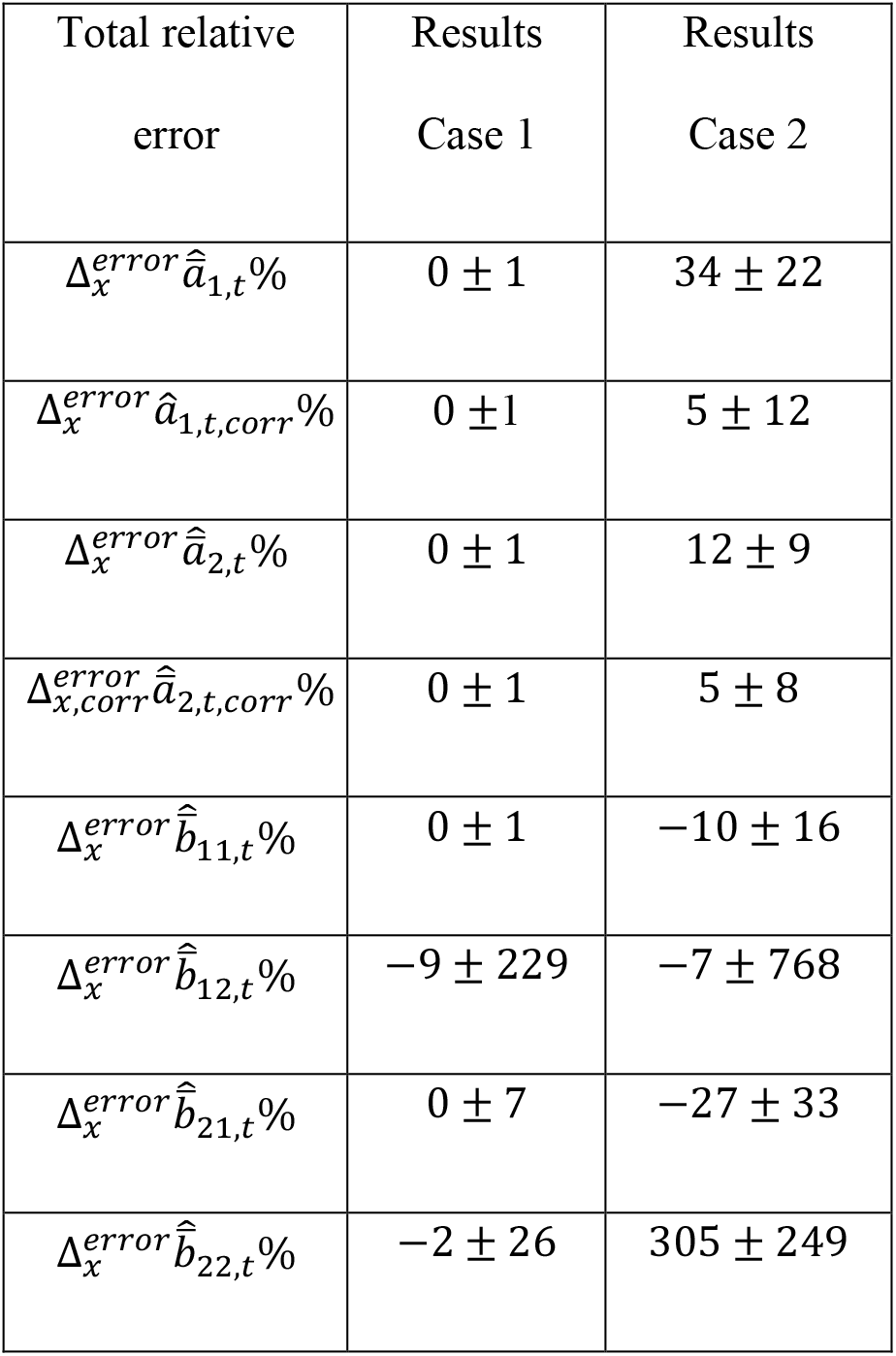
Mean trait prediction results with true system responses corresponding to Table 1.

As shown in Table 1, use of long input-output data from t=1 to *t* = 60 (Case 1) gives fairly good parameter estimates. The mean trait prediction results in Case 1 are correspondingly good, except for 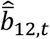 and 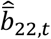 (Table 2). It is apparently difficult to separate the effects of the two environmental inputs, but a reduced model with 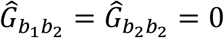 gave no improvement in other prediction results.

Use of short input-output data from t=31 to *t* = 60 (Case 2), and thus errors in 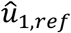 and 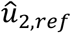, gives systematic errors in 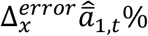 and 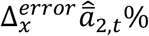, as discussed in Subsection 2.2. As shown in Table 2, these errors may be reduced by use of corrections as also discussed in Subsection 2.2, and the corrected results are clearly improved. There are large errors in 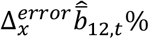 and 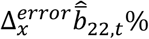, but also here a reduced model with 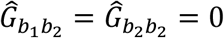 gave no improvement in other prediction results.

Fig. 4 shows typical plots of predicted mean values, as compared to true mean values for Case 1 in Table 1, i.e., with use of long data.

**Figure 4.**
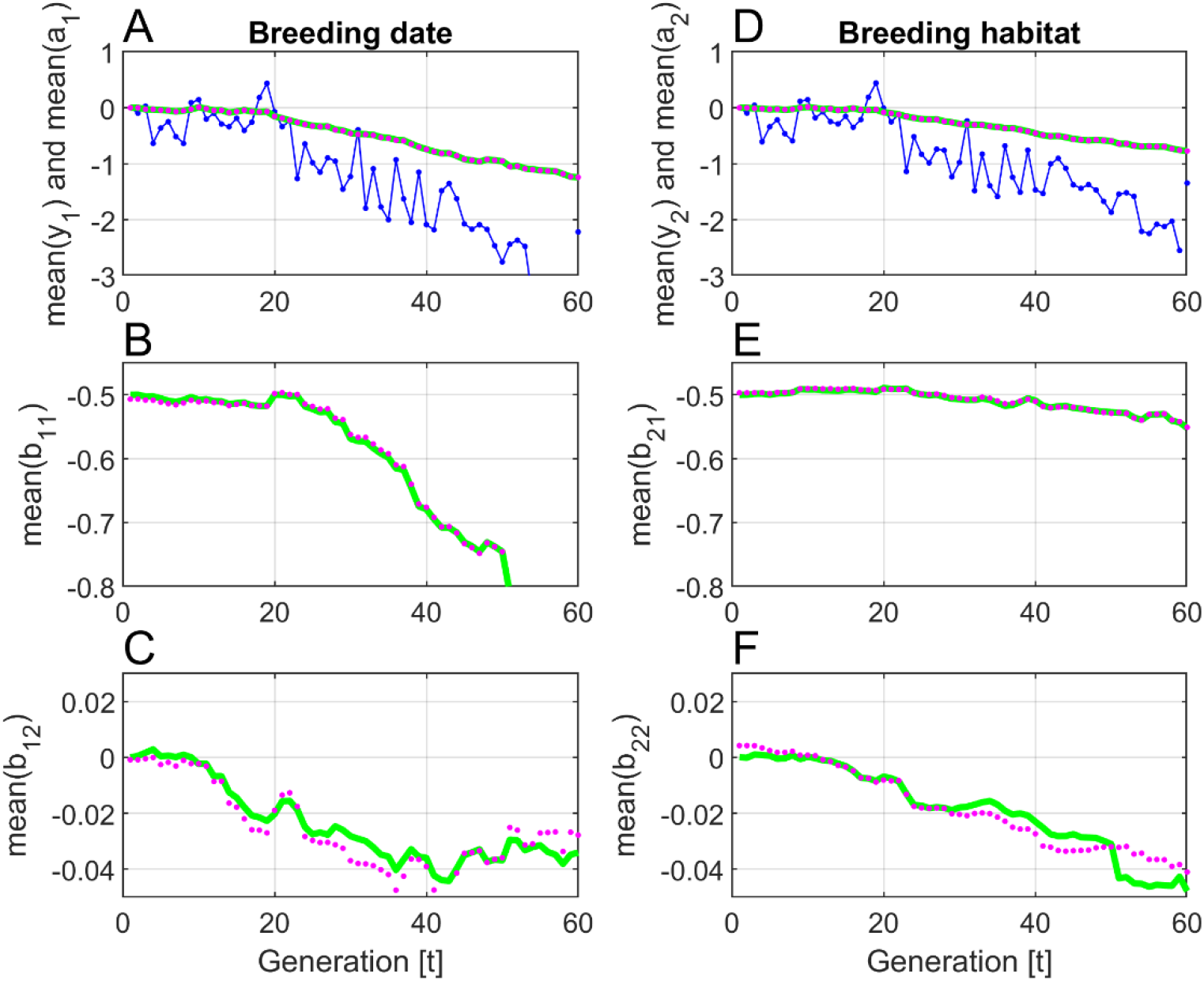
Typical responses for Case 1 in Table 1. All parameter values except 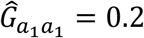 were initially set to zero. True 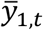 and 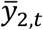 values are shown by solid blue lines, while predictions 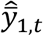 and 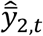 are shown by blue dots. True mean trait responses are shown by green lines, while predictions are shown by magenta dots.

Fig. 5 shows plots of predicted mean values, as compared to true mean values for Case 2 in Table 1.

**Figure 5.**
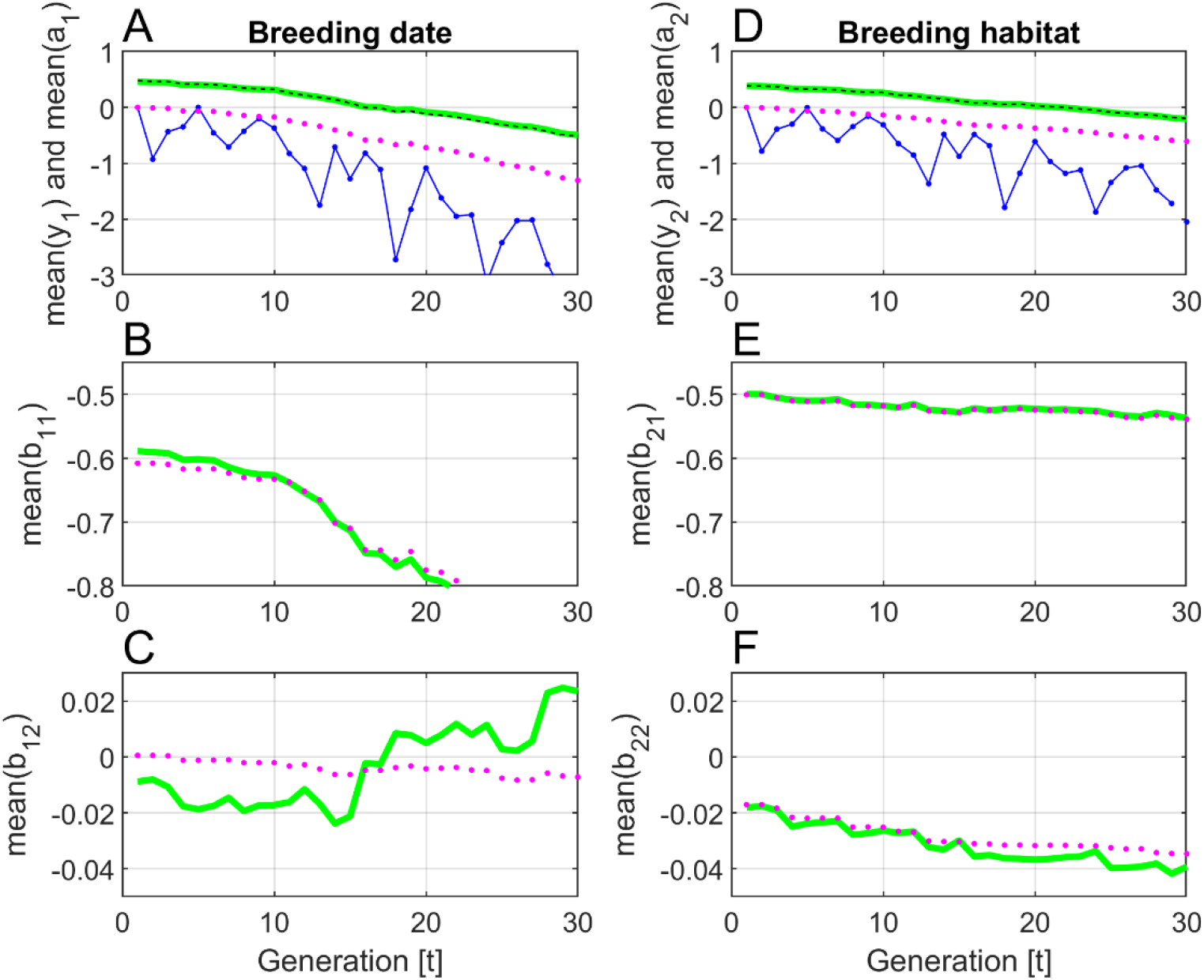
Typical responses for Case 2 in Table 1, although especially the plots in Panel C varies a lot from realization to realization. All parameter values except 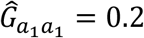 were initially set to zero. True 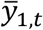 and 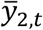 values are shown by solid blue lines, while final predictions 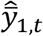 and 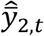 are shown by blue dots. True mean trait responses are shown by green lines, while predictions are shown by magenta dots. Corrected predictions as discussed in Subsection 2.2 are shown by black dashed lines.

## 4 Discussion and conclusions

The microevolutionary system identification method introduced in Ergon (2022) is here generalized from SISO to MIMO systems. The prediction error method (PEM) in Fig. 1 (with obvious notational changes for MIMO systems) is tested by simulations of a toy system with two environmental input signals (temperature and rainfall, Fig. 2), and two adaptive phenotypic output signals (mean breeding date and mean habitat choice). This system does not necessarily have realistic structure and parameter values, but the simulations still show that PEM works well for a complex system, provided that long input-output data that includes the environmental reference values are available (Table 1, Case 1, and Fig. 4). With short data that do not include the reference values, however, severe convergence problems are encountered, which is hardly surprising with in all 16 parameter values to be estimated (Fig. 3). Such problems may be solved by use of the first input values in the available time series (or similar values) as reference values (Table 1, Case 2, and Fig. 5, although also in this case 14% of the realizations showed signs of poor convergence). This leads to prediction errors, but these errors can to some degree be corrected, provided that the true reference values are approximately known (Table 2).

The difficulties encountered with parameter estimation and mean trait predictions based on short data must be expected also when other methods are used, for example best linear unbiased predictions (BLUP) combined with restricted maximum likelihood (REML) estimation (Ch. 26 and 27, Lynch and Walsh, 1998; Arnold *et al*., 2019). Note that PEM has a close kinship with maximum likelihood methods (Ljung, 2002).

A drawback with PEM as used in the simulations is that it assumes that there is no inbreeding in the population, i.e., that the relationship matrix is a unity matrix. However, as shown in a manuscript under review, it is possible to apply the PEM method also on BLUP models, where the relationship matrix has non-diagonal elements different from zero.

Another drawback with the theoretical treatment is that it assumes non-overlapping generations. There are, however, straightforward solutions for use of the multivariate breeder’s equation in cases with overlapping generations (Lande, 1979; Ch. 13, Walsh and Lynch, 2018). Since the prediction equations (6a,b) are derived by use of the breeder’s equation (Ergon, 2022), it should therefore be straightforward to find solutions for predictions with overlapping generations, but the details remain to be worked out.

## Supporting information

MATLAB code

## Acknowledgement

I thank University of South-Eastern Norway for support.

## Author Contribution

Rolf Ergon is the sole author of this article.

## Data Accessibility Statement

MATLAB code for simulations is given in Supplementary Material archived in *bioRxiv*, doi ?????

## References

Arnold, P.A., Kruuk, L.E.B., and Nicotra, A.B. (2019). How to analyse plant phenotypic plasticity in response to a changing climate. New Phytologist 222, 1235–1241. doi: 10.1111/nph.15656

Bowers, E.K., Grindstaff, J.L., Soukup, S.S., Drilling, N.E., Eckerle, K.P., Sakaluk, S.K., and Thompson, C.F. (2016). Spring temperatures influence selection on breeding date and the potential for phenological mismatch in a migratory bird. Ecology, 97(10), 2880–2891.

Chalfoun, A.D., and Schmidt, K.A. (2012). Adaptive breeding-habitat selection: Is it for the birds? The Auk, 129 (4), 589–599. doi: 10.1525/auk.2012.129.4.589

Ergon, T., and Ergon, R. (2017). When three traits make a line: evolution of phenotypic plasticity and genetic assimilation through linear reaction norms in stochastic environments. J. Evol. Biol. 30, 486–500. doi: 10.1111/jeb.13003

Ergon, R. (2019). Quantitative genetics state-space modeling of phenotypic plasticity and Evolution. Modeling, Identification and Control. 40, 51–69. Doi: 10.4173/mic.2019.1.5

Ergon, R. (2022). The important choice of reference environment in microevolutionary climate response predictions. Ecology and Evolution. doi: 10.1002/ece3.8836

Lande, R. (1979). Quantitative genetic analysis of multivariate evolution, applied to brain:body size allometry. Evolution 33, 402–416.

Lande, R. (2009). Adaptation to an extraordinary environment by evolution of phenotypic plasticity and genetic assimilation. J. Evol. Biol. 22, 1435–1446. doi: 10.1111/j.1420-9101.2009.01754.x

Ljung, L. (2002). Prediction Error Estimation Methods. Circuits, Systems, and Signal Processing 21/1, 11–21.

Lynch, M., and Walsh, B. (1998). Genetics and Analysis of Quantitatve Traits. Sinauer Associates, Mass.

Merilä, J., and Hendry, A.P. (2013). Climate change, adaptation, and phenotypic plasticity: the problem and the evidence. Evol. Appl. 7, 1–14. doi: 10.1111/eva.12137.

NASA Goddard Institute for Space Studies. (2019). Global Temperature Time Series. https://datahub.io/core/global-temp

Shaw, R.G., and Etterson, J.R. (2012). Rapid climate change and the rate of adaptation: insight from experimental quantitative genetics. New Phytologist 195: 752–765 doi: 10.1111/j.1469-8137.2012.04230.x

Walsh, B., and Lynch, M. (2018). _Evolution and Selection of Quantitative Traits. Oxford University Press

