## Supplementary material for "Microevolutionary system identification and climate response predictions": MATLAB code

### MATLAB code for

### Microevolutionary system identification

### and climate response predictions


Rolf Ergon

University of South-Eastern Norway


```
%----------------------------------------------------------
% Parameter estimation and mean trait prediction for MIMO model
% with use of fmincon in MATLAB.

% Rolf Ergon
% University of South-Eastern Norway
% April 23, 2022
%----------------------------------------------------------

clear
count=0;      % Count realiztions with fval<0.025

for m=1:100
    m
T=60;
x=60;         % x=60 (Case 1) or x=30 (Case 2)
N=400;
wsquare=10;
vartheta=2;
varu=0.5;
rho=0.25;
Gaa11=0.2;
Gaa22=0.2;
Gaa12=0.1;
Gbb11=0.05;
Gbb12=0.05;
Gbb21=0.05;
Gbb22=0.05;

Gaa=[Gaa11 Gaa12 ; Gaa12 Gaa22];
Gbb=diag([Gbb11 Gbb12 Gbb21 Gbb22]);
Gab=zeros(2,4);

varv11=Gaa11;
varv22=Gaa22;
vareta11=Gbb11;
vareta12=Gbb12;
vareta21=Gbb21;
vareta22=Gbb22;
Paa=[Gaa11+varv11 Gaa12 ; Gaa12 Gaa22+varv22];
Pbb=diag([Gbb11+vareta11 Gbb12+vareta12 Gbb21+vareta21 Gbb22+vareta22]);
G0=[Gaa Gab ; Gab' Gbb];
P0=[Paa Gab ; Gab' Pbb];

%% Generate u and theta sequences
uplot=zeros(1,T);
du1=zeros(1,T);
du2=zeros(1,T);
```

```
dtheta=zeros(1,T);
for i=2:T
    du1(i)=sqrt(varu)*randn;
    dtheta1(i)=du1(i)*rho*sqrt(vartheta/(varu))+sqrt(vartheta*(1-rho^2))*randn;
    if i>10
        uplot(i)=(i-10)/20;
    end
    du2(i)=0.5*du1(i)+0.5*sqrt(varu)*randn;
end
u10=uplot+du1;
u20=du2;
theta1=-2*uplot-dtheta1;
theta2=-1*uplot-du2;

%% Individual population traits around abar, bbar1 and bbar2
for t=1:T
    a1(:,t)=sqrt(Gaa11)*randn(N,1);
    a2(:,t)=Gaa12*a1(:,t)/Gaa11+sqrt(Gaa22-Gaa12^2/Gaa11)*randn(N,1);
    b11(:,t)=sqrt(Gbb11)*randn(N,1);
    b12(:,t)=sqrt(Gbb12)*randn(N,1);
    b21(:,t)=sqrt(Gbb21)*randn(N,1);
    b22(:,t)=sqrt(Gbb22)*randn(N,1);
    v1(:,t)=sqrt(varv11)*randn(N,1);
    v2(:,t)=sqrt(varv22)*randn(N,1);
    eta11(:,t)=sqrt(vareta11)*randn(N,1);
    eta12(:,t)=sqrt(vareta12)*randn(N,1);
    eta21(:,t)=sqrt(vareta21)*randn(N,1);
    eta22(:,t)=sqrt(vareta22)*randn(N,1);
    a1(:,t)=a1(:,t)-mean(a1(:,t));
    a2(:,t)=a2(:,t)-mean(a2(:,t));
    b11(:,t)=b11(:,t)-mean(b11(:,t));
    b12(:,t)=b12(:,t)-mean(b12(:,t));
    b21(:,t)=b21(:,t)-mean(b21(:,t));
    b22(:,t)=b22(:,t)-mean(b22(:,t));
    v1(:,t)=v1(:,t)-mean(v1(:,t));
    v2(:,t)=v2(:,t)-mean(v2(:,t));
    eta11(:,t)=eta11(:,t)-mean(eta11(:,t));
    eta12(:,t)=eta12(:,t)-mean(eta12(:,t));
    eta21(:,t)=eta21(:,t)-mean(eta21(:,t));
    eta22(:,t)=eta22(:,t) -mean(eta22(:,t));
end

%% Simulation of true system
abar10=0*ones(1,T);
abar20=0*ones(1,T);
bbar110=-0.5*ones(1,T);
bbar120=0*ones(1,T);
bbar210=-0.5*ones(1,T);
bbar220=0*ones(1,T);
ybar10=0*ones(1,T);
ybar20=0*ones(1,T);
for t=1:T-1
    bterm1=(bbar110(t)+b11(:,t)+eta11(:,t)).*u10(t)+(bbar120(t)+b12(:,t)+eta12(:,t)).*u20(t);
    bterm2=(bbar210(t)+b21(:,t)+eta21(:,t)).*u10(t)+(bbar220(t)+b22(:,t)+eta22(:,t)).*u20(t);
    Y10(:,t)=abar10(t)+a1(:,t)+v1(:,t)+bterm1;
    Y10(:,t)=round(10*Y10(:,t));
    Y10(:,t)=Y10(:,t)/10;
    Y20(:,t)=abar20(t)+a2(:,t)+v2(:,t)+bterm2;
    W0(:,t)=round(10*exp(-((Y10(:,t)-theta1(t)).^2+(Y20(:,t)-theta2(t)).^2)/(2*wsquare)));
    Wbar0(t)=mean(W0(:,t));
    covabW1=(N-1)*cov([a1(:,t)+v1(:,t) b11(:,t)+eta11(:,t) b12(:,t)+eta12(:,t) W0(:,t)])/N;
    covabW2=(N-1)*cov([a2(:,t)+v2(:,t) b21(:,t)+eta21(:,t) b22(:,t)+eta22(:,t) W0(:,t)])/N;
    xbar0(:,t)=[abar10(t) abar20(t) bbar110(t) bbar120(t) bbar210(t) bbar220(t)]';
    covxW=[covabW1(1,4) covabW2(1,4) covabW1(2,4) covabW1(3,4) covabW2(2,4) covabW2(3,4)];
    xbar0(:,t+1)=xbar0(:,t)+G0*inv(P0)*covxW'/Wbar0(t);
    xbarny=xbar0(:,t+1);
    abar10(t+1)=xbarny(1);
    abar20(t+1)=xbarny(2);
    bbar110(t+1)=xbarny(3);
    bbar120(t+1)=xbarny(4);
    bbar210(t+1)=xbarny(5);
```

```
    bbar220(t+1)=xbarny(6);
    ybar10(t+1)=abar10(t+1)+bbar110(t+1)*u10(t+1)+bbar120(t+1)*u20(t+1);
    ybar20(t+1)=abar20(t+1)+bbar210(t+1)*u10(t+1)+bbar220(t+1)*u20(t+1);
end

%% Short data
u1=u10(T-x+1:T)-u10(T-x+1);
u2=u20(T-x+1:T)-u20(T-x+1);
% u1=u10(T-x+1:T);
% u2=u20(T-x+1:T);
Y1=Y10(:,T-x+1:T-1);
Y2=Y20(:,T-x+1:T-1);
abar1=abar10(T-x+1:T)-ybar10(T-x+1);
abar2=abar20(T-x+1:T)-ybar20(T-x+1);
bbar11=bbar110(T-x+1:T);
bbar12=bbar120(T-x+1:T);
bbar21=bbar210(T-x+1:T);
bbar22=bbar220(T-x+1:T);
ybar1=ybar10(T-x+1:T-1)-ybar10(T-x+1);
ybar2=ybar20(T-x+1:T-1)-ybar20(T-x+1);
W=W0(:,T-x+1:T-1);
Wbar=mean(W);

%% Contraints
Gaa22_min=0;Gaa22_max=0.5;
Gaa12_min=-0.1;Gaa12_max=0.5;
Gbb11_min=0; Gbb11_max=0.2;
Gbb12_min=0; Gbb12_max=0.2;
Gbb21_min=0; Gbb21_max=0.2;
Gbb22_min=0; Gbb22_max=0.2;
varv11_min=0;varv11_max=1;
varv22_min=0;varv22_max=1;
vareta11_min=0;vareta11_max=0.2;
vareta12_min=0;vareta12_max=0.2;
vareta21_min=0;vareta21_max=0.2;
vareta22_min=0;vareta22_max=0.2;
bbarinit11_min=-1; bbarinit11_max=0;
bbarinit12_min=-1; bbarinit12_max=1;
bbarinit21_min=-1; bbarinit21_max=0;
bbarinit22_min=-1; bbarinit22_max=1;
Gaamin=[Gaa22_min,Gaa12_min];
Gaamax=[Gaa22_max,Gaa12_max];
Gbbmin=[Gbb11_min,Gbb12_min,Gbb21_min,Gbb22_min];
Gbbmax=[Gbb11_max,Gbb12_max,Gbb21_max,Gbb22_max];
varvmin=[varv11_min,varv22_min];
varvmax=[varv11_max,varv22_max];
varetamin=[vareta11_min,vareta12_min,vareta21_min,vareta22_min];
varetamax=[vareta11_max,vareta12_max,vareta21_max,vareta22_max];
bbarinitmin=[bbarinit11_min,bbarinit12_min,bbarinit21_min,bbarinit22_min];
bbarinitmax=[bbarinit11_max,bbarinit12_max,bbarinit21_max,bbarinit22_max];
par_lb=[Gaamin,Gbbmin,varvmin,varetamin,bbarinitmin];
par_ub=[Gaamax,Gbbmax,varvmax,varetamax,bbarinitmax];
par_guess=zeros(1,16);
par_known=[Gaa11];

%% fmincon
Aineq=[]; Bineq=[]; Aeq=[]; Beq=[];
fun_objective_MIMO_handle=...
    @(par)fun_objective_MIMO(par,par_known,u1,u2,Y1,Y2,ybar1,ybar2,W,Wbar,N,x);
fun_constraints_MIMO_handle=...
    @(par)fun_constraints_MIMO(par,par_known,u1,u2,Y1,Y2,ybar1,ybar2,W,Wbar,N,x);
[par_opt,fval,exitflag,output,lambda,grad,hessian] =...
fmincon(fun_objective_MIMO_handle,par_guess,Aineq,Bineq,Aeq,Beq,par_lb,par_ub,fun_constraints_MIMO_handle);
output
fval
Fval(m)=fval;
opt_results(m,:)=par_opt;

%% Simulation with adapted model
Gaa22=par_opt(1);
Gaa12=par_opt(2);
```

```
Gbb11=par_opt(3);
Gbb12=par_opt(4);
Gbb21=par_opt(5);
Gbb22=par_opt(6);
varv11=par_opt(7);
varv22=par_opt(8);
vareta11=par_opt(9);
vareta12=par_opt(10);
vareta21=par_opt(11);
vareta22=par_opt(12);
bbarinit11=par_opt(13);
bbarinit12=par_opt(14);
bbarinit21=par_opt(15);
bbarinit22=par_opt(16);
Gaa=[Gaa11 Gaa12 ; Gaa12 Gaa22];
Gbb=diag([Gbb11 Gbb12 Gbb21 Gbb22]);
Gab=zeros(2,4);
Paa=[Gaa11+varv11 Gaa12 ; Gaa12 Gaa22+varv22];
Pbb=diag([Gbb11+vareta11 Gbb12+vareta12 Gbb21+vareta21 Gbb22+vareta22]);
G=[Gaa Gab ; Gab' Gbb];
P=[Paa Gab ; Gab' Pbb];

abarhat=zeros(2,x);
bbarhat=[bbarinit11*ones(1,x)
        bbarinit12*ones(1,x)
        bbarinit21*ones(1,x)
        bbarinit22*ones(1,x)];
ybarhat1=ybar1(1)*ones(1,55);
ybarhat2=ybar2(1)*ones(1,55);
ybarhat=[ybarhat1;ybarhat2];

for t=1:x-1
   U=[u1(t) 0 ; u2(t) 0 ; 0 u1(t) ; 0 u2(t)];
   Pyy=Paa+U'*Pbb*U;
   covWy1=(N-1)*cov(W(:,t),Y1(:,t))/N;
   covWy2=(N-1)*cov(W(:,t),Y2(:,t))/N;
   betay(:,t)=inv(Pyy)*[covWy1(1,2) ; covWy2(1,2)]/Wbar(t);
   abarhat(:,t+1)=abarhat(:,t)+Gaa*betay(:,t);
   bbarhat(:,t+1)=bbarhat(:,t)+Gbb*U*betay(:,t);
   U=[u1(t+1) 0 ; u2(t+1) 0 ; 0 u1(t+1) ; 0 u2(t+1)];
   ybarhat(:,t+1)=abarhat(:,t+1)+U'*bbarhat(:,t+1);
   abarhat1(:,t+1)=abarhat(1,t+1);
   abarhat2(:,t+1)=abarhat(2,t+1);
   bbarhat11(:,t+1)=bbarhat(1,t+1);
   bbarhat12(:,t+1)=bbarhat(2,t+1);
   bbarhat21(:,t+1)=bbarhat(3,t+1);
   bbarhat22(:,t+1)=bbarhat(4,t+1);
end
abarhat1=abarhat(1,:);
abarhat2=abarhat(2,:);
bbarhat11=bbarhat(1,:);
bbarhat12=bbarhat(2,:);
bbarhat21=bbarhat(3,:);
bbarhat22=bbarhat(4,:);
ybarhat1=ybarhat(1,:);
ybarhat2=ybarhat(2,:);

abarhat1corr=abarhat1-bbarhat11*(u10(T-x+1))-bbarhat12*(u20(T-x+1));
abarhat2corr=abarhat2-bbarhat21*(u10(T-x+1))-bbarhat22*(u20(T-x+1));

if fval<0.025
   count=count+1;
   Totalabar1=abar1(1)-abar1(x);
   Totalabarhat1=abarhat1(1)-abarhat1(x);
   Rel_total_abar_error_1(count)=(Totalabarhat1-Totalabar1)/Totalabar1;
   Totalabarhat1corr=abarhat1corr(1)-abarhat1corr(x);
   Rel_total_abar_error_1_corr(count)=(Totalabarhat1corr-Totalabar1)/Totalabar1;

   Totalabar2=abar2(1)-abar2(x);
   Totalabarhat2=abarhat2(1)-abarhat2(x);
   Rel_total_abar_error_2(count)=(Totalabarhat2-Totalabar2)/Totalabar2;
```

```
    Totalabarhat2corr=abarhat2corr(1)-abarhat2corr(x);
    Rel_total_abar_error_2_corr(count)=(Totalabarhat2corr-Totalabar2)/Totalabar2;

    Totalbbar11=bbar11(1)-bbar11(x);
    Totalbbarhat11=bbarhat11(1)-bbarhat11(x);
    Rel_total_bbar_error_11(count)=(Totalbbarhat11-Totalbbar11)/Totalbbar11;

    Totalbbar12=bbar12(1)-bbar12(x);
    Totalbbarhat12=bbarhat12(1)-bbarhat12(x);
    Rel_total_bbar_error_12(count)=(Totalbbarhat12-Totalbbar12)/Totalbbar12;

    Totalbbar21=bbar21(1)-bbar21(x);
    Totalbbarhat21=bbarhat21(1)-bbarhat21(x);
    Rel_total_bbar_error_21(count)=(Totalbbarhat21-Totalbbar21)/Totalbbar21;

    Totalbbar22=bbar22(1)-bbar22(x);
    Totalbbarhat22=bbarhat22(1)-bbarhat22(x);
    Rel_total_bbar_error_22(count)=(Totalbbarhat22-Totalbbar22)/Totalbbar22;
end

end

%% Plots:

figure(4)
subplot(3,2,1)
plot(ybar1,'b'), hold on
plot(ybarhat1,'.b','LineWidth',2)
plot(abar1,'g','LineWidth',2)
plot(abarhat1,'.m','LineWidth',2)
plot(abarhat1corr,'--k')
title('Breeding date')
ylabel('mean(y_1) and mean(a_1)')
axis([0 x -3 1]), hold off, grid
text(0,1.5,'A','FontSize',14)

subplot(3,2,3)
plot(bbar11,'g','LineWidth',2), hold on
plot(bbarhat11,'.m','LineWidth',2)
% plot(bbar1e,'.b'),
hold off, grid
ylabel('mean(b_1_1)')
axis([0 x -0.8 -0.45])
text(0,-0.41,'B','FontSize',14)

subplot(3,2,5)
plot(bbar12,'g','LineWidth',2), hold on
plot(bbarhat12,'.m','LineWidth',2)
% plot(bbar2e,'.b'),
hold off, grid
xlabel('Generation [t]')
ylabel('mean(b_1_2)')
axis([0 x -0.05 0.03])
text(0,0.04,'C','FontSize',14)

subplot(3,2,2)
plot(ybar2,'b'), hold on
plot(ybarhat2,'.b','LineWidth',2)
plot(abar2,'g','LineWidth',2)
plot(abarhat2,'.m','LineWidth',2)
plot(abarhat2corr,'--k')
title('Breeding habitat')
ylabel('mean(y_2) and mean(a_2)')
axis([0 x -3 1])
hold off
grid
text(0,1.5,'D','FontSize',14)

subplot(3,2,4)
plot(bbar21,'g','LineWidth',2), hold on
plot(bbarhat21,'.m','LineWidth',2)
```

```
% plot(bbar1e,'.b'),
hold off, grid
ylabel('mean(b_2_1)')
axis([0 x -0.8 -0.45])
text(0,-0.41,'E','FontSize',14)

subplot(3,2,6)
plot(bbar22,'g','LineWidth',2), hold on
plot(bbarhat22,'.m','LineWidth',2)
% plot(bbar2e,'.b'),
hold off, grid
xlabel('Generation [t]')
ylabel('mean(b_2_2)')
axis([0 x -0.05 0.03])
text(0,0.04,'F','FontSize',14)

%% Results
Mean=mean(opt_results)
Std=std(opt_results)

Mean_abar_error_1=mean(Rel_total_abar_error_1);
Std_abar_error_1=std(Rel_total_abar_error_1);
abar1_results=[Mean_abar_error_1 Std_abar_error_1]

Mean_abar_error_1_corr=mean(Rel_total_abar_error_1_corr);
Std_abar_error_1_corr=std(Rel_total_abar_error_1_corr);
abar1_results_corr=[Mean_abar_error_1_corr Std_abar_error_1_corr]

Mean_abar_error_2=mean(Rel_total_abar_error_2);
Std_abar_error_2=std(Rel_total_abar_error_2);
abar2_results=[Mean_abar_error_2 Std_abar_error_2]

Mean_abar_error_2_corr=mean(Rel_total_abar_error_2_corr);
Std_abar_error_2_corr=std(Rel_total_abar_error_2_corr);
abar2_results_corr=[Mean_abar_error_2_corr Std_abar_error_2_corr]

Mean_bbar_error_11=mean(Rel_total_bbar_error_11);
Std_bbar_error_11=std(Rel_total_bbar_error_11);
bbar11_results=[Mean_bbar_error_11 Std_bbar_error_11]

Mean_bbar_error_12=mean(Rel_total_bbar_error_12);
Std_bbar_error_12=std(Rel_total_bbar_error_12);
bbar12_results=[Mean_bbar_error_12 Std_bbar_error_12]

Mean_bbar_error_21=mean(Rel_total_bbar_error_21);
Std_bbar_error_21=std(Rel_total_bbar_error_21);
bbar21_results=[Mean_bbar_error_21 Std_bbar_error_21]

Mean_bbar_error_22=mean(Rel_total_bbar_error_22);
Std_bbar_error_22=std(Rel_total_bbar_error_22);
bbar22_results=[Mean_bbar_error_22 Std_bbar_error_22]


for t=1:30
   bsum1(t)=bbar11(t)*u1(t)+bbar12(t)*u2(t);
   bsum2(t)=bbar21(t)*u1(t)+bbar22(t)*u2(t);
   bsumhat1(t)=bbarhat11(t)*u1(t)+bbarhat12(t)*u2(t);
   bsumhat2(t)=bbarhat21(t)*u1(t)+bbarhat22(t)*u2(t);
end

figure(2)
u2plot=2*ones(1,60);
subplot(2,2,1)
plot(u10+10), hold on
plot(10+uplot,'--'), hold off
axis([0 60 8 13]), grid
ylabel('u_1 (temperature)')
text(0,13.4,'A','FontSize',14)
subplot(2,2,2)
plot(u20+2), hold on
plot(u2plot,'--'), hold off
```

```
axis([0 60 0 5]), grid
ylabel('u_2 (rainfall)')
text(0,5.4,'C','FontSize',14)
subplot(2,2,3)
plot(theta1), hold on
plot(-2*uplot,'--'), hold off
axis([0 60 -6 2]), grid
xlabel('Generation [t]')
ylabel('theta_1')
text(0,2.6,'B','FontSize',14)
subplot(2,2,4)
plot(theta2), hold on
plot(-uplot,'--'), hold off
axis([0 60 -6 2]), grid
xlabel('Generation [t]')
ylabel('theta_2')
text(0,2.6,'D','FontSize',14)

count

%% Objective function
function f = fun_objective_MIMO(par,par_known,u1,u2,Y1,Y2,ybar1,ybar2,W,Wbar,N,x)
Gaa11=par_known;
Gaa22=par(1);
Gaa12=par(2);
Gbb11=par(3);
Gbb12=par(4);
Gbb21=par(5);
Gbb22=par(6);
varv11=par(7);
varv22=par(8);
vareta11=par(9);
vareta12=par(10);
vareta21=par(11);
vareta22=par(12);
bbarinit11=par(13);
bbarinit12=par(14);
bbarinit21=par(15);
bbarinit22=par(16);
Gaa=[Gaa11 Gaa12 ; Gaa12 Gaa22];
Gbb=diag([Gbb11 Gbb12 Gbb21 Gbb22]);
Gab=[0 0 0 0
    0 0 0 0];
Paa=[Gaa11+varv11 Gaa12 ; Gaa12 Gaa22+varv22];
Pbb=diag([Gbb11+vareta11 Gbb12+vareta12 Gbb21+vareta21 Gbb22+vareta22]);
G=[Gaa Gab ; Gab' Gbb];
P=[Paa Gab ; Gab' Pbb];
abarhat=zeros(2,x);
bbarhat=[bbarinit11*ones(1,x)
       bbarinit12*ones(1,x)
       bbarinit21*ones(1,x)
       bbarinit22*ones(1,x)];
ybaropj=zeros(2,x-1);

for t=1:x-1
   U=[u1(t) 0 ; u2(t) 0 ; 0 u1(t) ; 0 u2(t)];
   Pyy=Paa+U'*Pbb*U;
   covWy1=(N-1)*cov(W(:,t),Y1(:,t))/N;
   covWy2=(N-1)*cov(W(:,t),Y2(:,t))/N;
   betay(:,t)=inv(Pyy)*[covWy1(1,2) ; covWy2(1,2)]/Wbar(t);
   abarhat(:,t+1)=abarhat(:,t)+Gaa*betay(:,t);
   bbarhat(:,t+1)=bbarhat(:,t)+Gbb*U*betay(:,t);
   U=[u1(t+1) 0 ; u2(t+1) 0 ; 0 u1(t+1) ; 0 u2(t+1)];
   ybarobj(:,t+1)=abarhat(:,t+1)+U'*bbarhat(:,t+1);
end
ybarobj1=ybarobj(1,:);
ybarobj2=ybarobj(2,:);
f=sum(0.5*(ybar1(1:x-1)-ybarobj1(1:x-1)).^2)+sum(0.5*(ybar2(1:x-1)-ybarobj2(1:x-1)).^2);

end
```

```
%% Constraints function
function [cineq,ceq]=fun_constraints_MIMO(par,par_known,u1,u2,Y1,Y2,ybar1,ybar2,W,Wbar,N,x)
cineq =  [];   % Compute nonlinear inequalities.
ceq = [];   % Compute nonlinear equalities.
end
```
